# Predicting complex genetic phenotypes using error propagation in weighted networks

**DOI:** 10.1101/487348

**Authors:** El Mahdi El Mhamdi, Andrei Kucharavy, Rachid Guerraoui, Rong Li

## Abstract

Network-biology view of biological systems is a ubiquitous abstraction that emerged in the last two decades to allow a high-level understanding of principles governing them. However, the principles according to which biological systems are organized are still unclear. Here, we investigate if biological networks could be approximated as overlapping, feed-forward networks where the nodes have non-linear activation functions. Such networks have been shown to be universal approximators and their stability has been explored in the context of artificial neural networks. Mathematical formalization of this model followed by numerical simulations based on genomic data allowed us to accurately predict the statistics of gene essentiality in yeast and hence indicate that biological networks might be better understood as a distributed system, comprising potentially unreliable components.

## Main Text

With the advent of the genome-wide association studies (GWAS), the biomedical community has become increasingly aware of and interested in complex genetic phenotypes. At the population level, diseases such as cancer (*1*), schizophrenia (*2*) or autism (*3*) are not associated with a small set of causative genes but are rather caused by a large number of perturbations distributed across numerous genes that lead to forms of disorders differing in severity and age of onset. Further investigation has shown that such complex phenotypes are not an exception but rather a rule. Survival and growth in rich medium upon genes deletion, a classical model for the study of the genetic basis of a phenotype (survival) has recently revealed itself to be a complex genetic phenotype. Genes required for survival—essential genes—are not defined in absolute terms, but can be dispensable in some environments or genetic contexts (4, 5). Understanding and predicting complex genetic phenotypes is a crucial step towards getting an actionable handle on biologically and clinically relevant phenomena.

Similarly to the previous work (*6*) a biological system is viewed as a directed weighted graph, where transcription factors and their regulated genes are nodes, and regulatory relationships are edges. We go one step further and recognize any metabolite, DNA or RNA fragment, and protein or complex as a node. We associate a node with chemical reactions producing it. We refer to this abstraction as BOWN, standing for Biological Organisms as Weighted Networks. Within BOWN, molecules are the nodes while edges describe involvement in biochemical reactions that change node states or transform them from one into another, weighted by coefficients encompassing stoichiometry and activation intensity. While a variety of activation functions are present in biological systems, we assume that they have minimum and maximum values and a finite slope between the two. A sigmoidal activation function is used because it simplifies the mathematical analysis and corresponds to an idealized form of the Hill equation in biochemistry, representing ultrasensitive saturating activation by one or more factors.

We applied our formalism to the understanding of essential genes, traditionally defined as genes whose deletion is lethal to the organism. Essential genes in unicellular organisms are usually highly conserved (*7*), hence considered as performing critical core functions (*8, 9*) and of prime importance as targets to new drugs against unicellular pathogens (*10*). In a well-studied model organism, the yeast *Saccharomyces Cerevisiae*, about 1100 of the total 6500 genes are essential for growth on rich media (*11, 12*). Recently, however, the essentiality of some of those genes has been questioned. *MYO1*, an essential gene in *S. Cerevisiae*, could be deleted without leading to death, given that the genome was perturbed at a large scale by chromosome copy number variation (*5*). The same work also showed that at least 3 different mechanisms could compensate for *MYO1* deletion. Among those mechanisms, none relied on paralogs. Later, a systemic genome-wide analysis showed that 9% of all essential genes behave similarly and termed such genes “evolvable essential” (*4*). These recent studies raised new questions on the origin of essential genes, such as how essential genes come to be and what separates them from non-essential ones.

In the past four decades, the theory of fault tolerance in distributed systems, a branch of computer science (*13–15*) that studies systems ability to tolerate the failure of some their components, has led to an abstraction to describe nodes whose loss leads to critical failure in a network as *single*^1^ *points of failure;* in this formalism, a necessary condition for a system to be robust is not to have any single point of failure. The more failing nodes can be tolerated, the more robust is the system. Based on this definition, biological networks are not robust with regards to survival as essential genes represent single points of failure. Computer scientists might note that biomolecular networks are not composed of redundant nodes performing the same task and trying to reach an agreement on the results. Hence, biomolecular networks do not conform to core assumptions that underlie most results in distributed computing. Rather than agreement-centered distributed systems, BOWN leverages a fault tolerance formalism that is reminiscent of the one used for neural networks (*16–18*) which like BOWN, can be seen as (layered) weighted directed graphs with non-linear nodes.

Within BOWN, each node’s state depends on some other nodes. Among those, some nodes are more important than others -we say nodes weight each other’s output differently. Most importantly, these nodes perform different tasks, despite some overlapping functions. Hence, none of the agreement-centered theories (*13–15*), traditional to distributed computing, can be directly used to understand the robustness of biomolecular networks. Yet, the very notion of a *weighted directed graph*, defined as a set of computing nodes, all interdependent for computation, is an abstraction that could be used to represent biological networks. This notion has already been applied in genetics context (*6*), and recently as an error propagation model in distributed computing systems such as biological neural networks (*19*) or artificial ones (*4, 16, 17*). We use this abstraction of weighted directed graphs with nonlinear nodes to model biomolecular networks. Here, non-linear nodes are nodes whose output is not necessarily in a linear relationship with their input. Sigmoidal functions, for instance occurring in reactions following Hill kinetics, are examples of such non-linear relationships between the inputs (reagents) and the outputs (products).

A distributed computing system has two main types of components, processes and communication channels (*13*). In the context of directed weighted graphs, processes are nodes and edges are channels. In the context of biomolecular networks, *processes* are proteins, RNA or small molecules identified with the reactions producing them. The reagents then route the information through a biological network by changing their state through biochemical processes, be it the modification of compartmental localization, participation in regulatory or enzymatic complexes, or the transformation into a different physical entity by chemical reaction. The biochemical relationships between those processes are viewed as the *communication channels*. The information is transmitted between nodes through the functional modification of one node by the other, be it by reaction co-participation, complex formation, or post-translational modification. Figure 1 shows the correspondences between those nodes in distributed computing (A), directed graphs (B) as well as potential nodes and edges in biological systems (C, D).

**Fig. 1.**
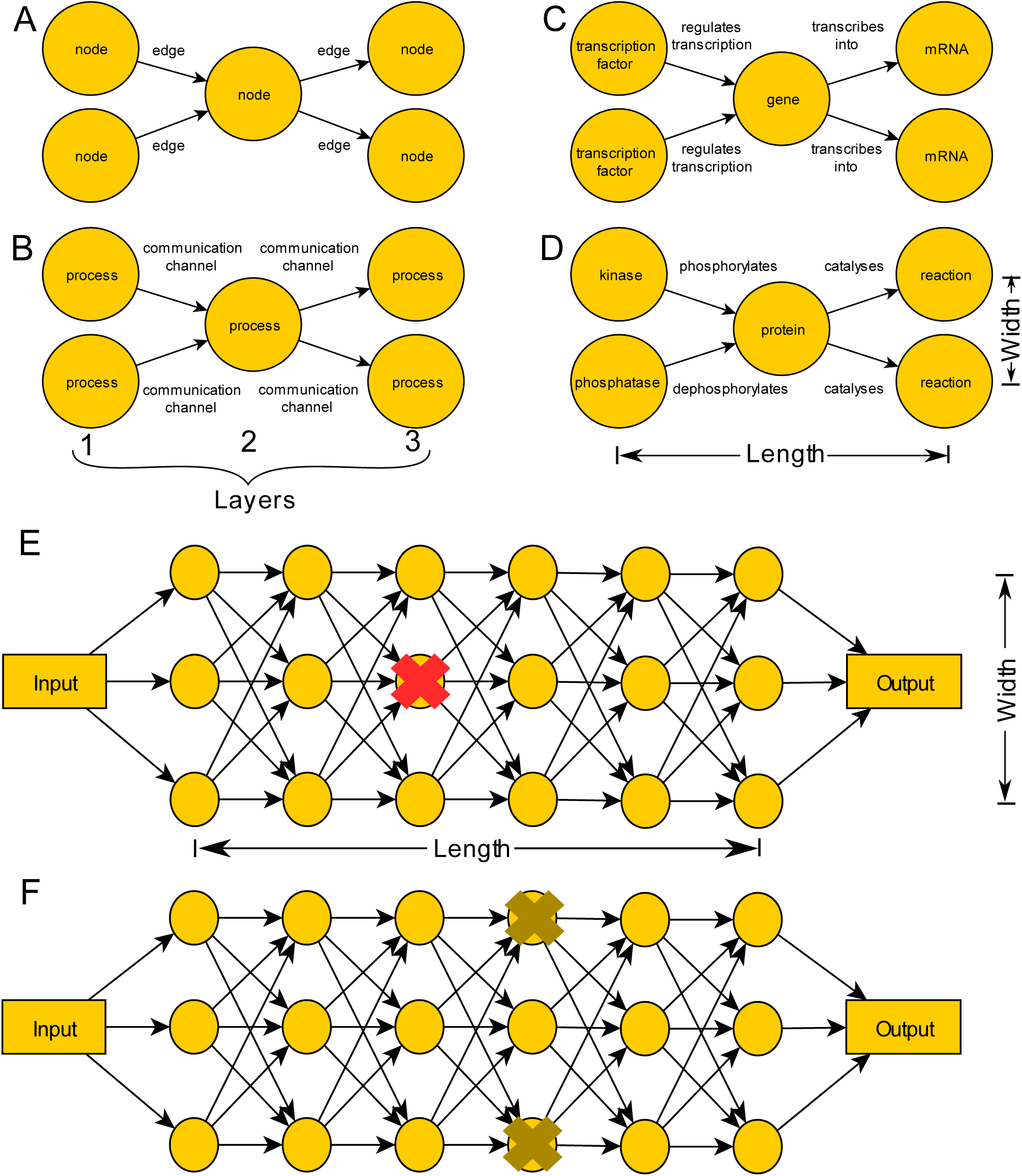
Examples of networks and BOWN network model (A) Directed graph. (B) Distributed computation. (C, D): Examples of biological implementations of a network. (A-D) All networks have three layers (length of 3) with per-layer width of up to two. (E) Maximum error propagation network. The error is calculated on the output, single deletions leading to over 70% output error correspond to essential genes (red cross). (F) Double deletions of non-essential genes leading to an error above 70% are considered as SLs (light brown)

We model a biomolecular network by a layered directed graph (BOWN) as depicted in Figure 1B. Weighted directed graphs with nonlinear nodes are proven to be universal computing objects (*20, 21*), as such they can map any environment (input) to any biological system response (output) given enough nodes. The most explored formalization of these systems are artificial neural networks (*22*). In BOWN, we approximate biological pathways by such layered networks (examples in Figure 1C-D). To analyze such network’s stability, we need to look at their properties. First, the reactions are thermodynamically stable, meaning that no change in concentration of a reagent will induce an unbounded change in the concentration of a reagent downstream. In the parlance of distributed computing, the transmission channel is limited in throughput. Second, we model reactions by the sigmoidal behavior described by a Hill equation. The sigmoidal behavior is not a requirement for BOWN – any function with lower and upper limit, and a continuous transition between the two is sufficient as proven previously (*20, 21*). These two considerations allow us to derive mathematical conditions on what it means to be *essential* in a weighted directed graph such as BOWN. This theoretical development is presented more in-depth in (*23*) (Lemma 1, Proposition 1 and Proposition 2) and provides us with the main formula for the identification of essential genes. Briefly, we denote by *l* the layer index, *i, j* identifiers of nodes in layers, *c_i,j_* the weight of the link between node *i* and node *j*, 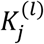 the Hill coefficient connecting nodes in layer *l* − 1 to node *j* in layer *l* and finally, *∊* the maximum tolerated error in the functional pathway (beyond this error, the organism dies). A gene *k* in layer *l′* is considered essential if the suppression of *k* yields a lethal value for the forward-propagated error at the output, which corresponds to the condition below:

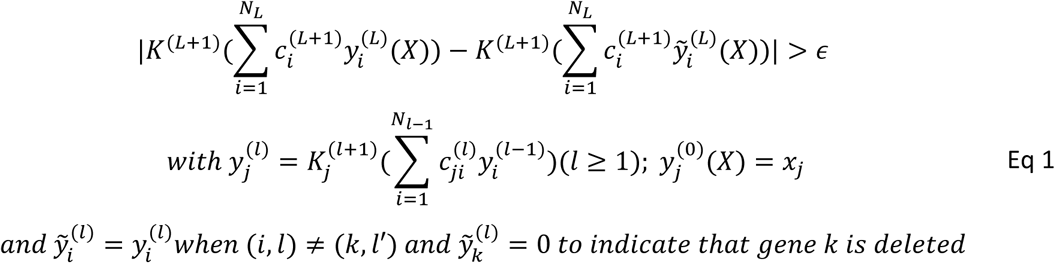

The inequality in Eq 1 describes the following: If a node disruption affects another node that is important or is a main regulator of sufficiently important proteins, this would lead to a large error in the response of the pathway to the input. The effect becomes lethal when this error exceeds *∊*, i.e the organism is not able to recover from the error propagation described by the left-hand side term (Figure 1E). Similarly, BOWN allows to check if a disruption of several nodes can lead to a similar lethal error (Figure 1F).

Equation 1 means that our model predicts that the node criticality, and hence the essentiality of the corresponding gene, is amplified by one (or a combination) of three factors. (*1*) High outgoing weights - essential genes are likely to be limiting factors in a number of biochemical reactions. (*2*) Strong links to downstream nodes with high kinetic factors (high 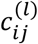 and 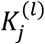 in Equation S4; since we are interested in maximum error, Equation S4 assumes the entire network is at the point of highest slope of activation function). (3) Being as far upstream as possible in the pathways, and everything else being equal. It is important to note that if the exact topology and weights of a biological network were provided, the same approach that led to Eq 1 – quantifying error propagation - would pinpoint the critical nodes exactly and hence allow us to identify specific essential genes by their indexes. Unfortunately, the availability of the weights connecting all the nodes are not known, and we hence focus on the prevalence of critical nodes in the network, without attempting to predict their identities. These types of predictions can be based on (1) statistics describing the weights connecting the nodes, (2) the distribution of lengths and widths of pathways, and (3) the maximal tolerated error *∊*.

Due to BOWN’s similarity with neural-networks models, we expect that the statistics of these three parameters reflect a trade-off that the organism has to strike between robustness, evolvability and network size - the latter being limited by resources - as demonstrated previously (*16,17*). Given that those constraints are general, we opted not to focus on specific networks, such as gene regulation, physical protein-protein interaction or phosphorylation network, but rather use combined statistics to characterize them.

First, we estimate a biologically plausible distribution of weights connecting one node to others based on the data from a transcription factors (TF) deletion profiling (*6*). Hu *et al*. measured expression levels of all genes in yeast upon deletion of a TF from a set of 250 TFs. Figure 2A represents the pooled distribution of gene expression changes for all TF deletions. We use this distribution to sample the multiplicative products of the kinetic factors by the stoichiometric ones (corresponding to 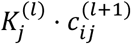 in Equation 2). While the effect of TF deletions on the vast majority of genes are within the margin of experimental error (red on Figure 2A), we can see at the extremes of the distribution of a small fraction of genes whose transcription is strongly regulated by a particular TF, directly or indirectly. Given the sparsity of connections within real biological pathways (most proteins tend to interact only with a few proteins and most transcription factors regulate expression of only a fraction of genes), we sampled the weights for our model from the whole distribution and not only from the extrema.

**Fig. 2.**
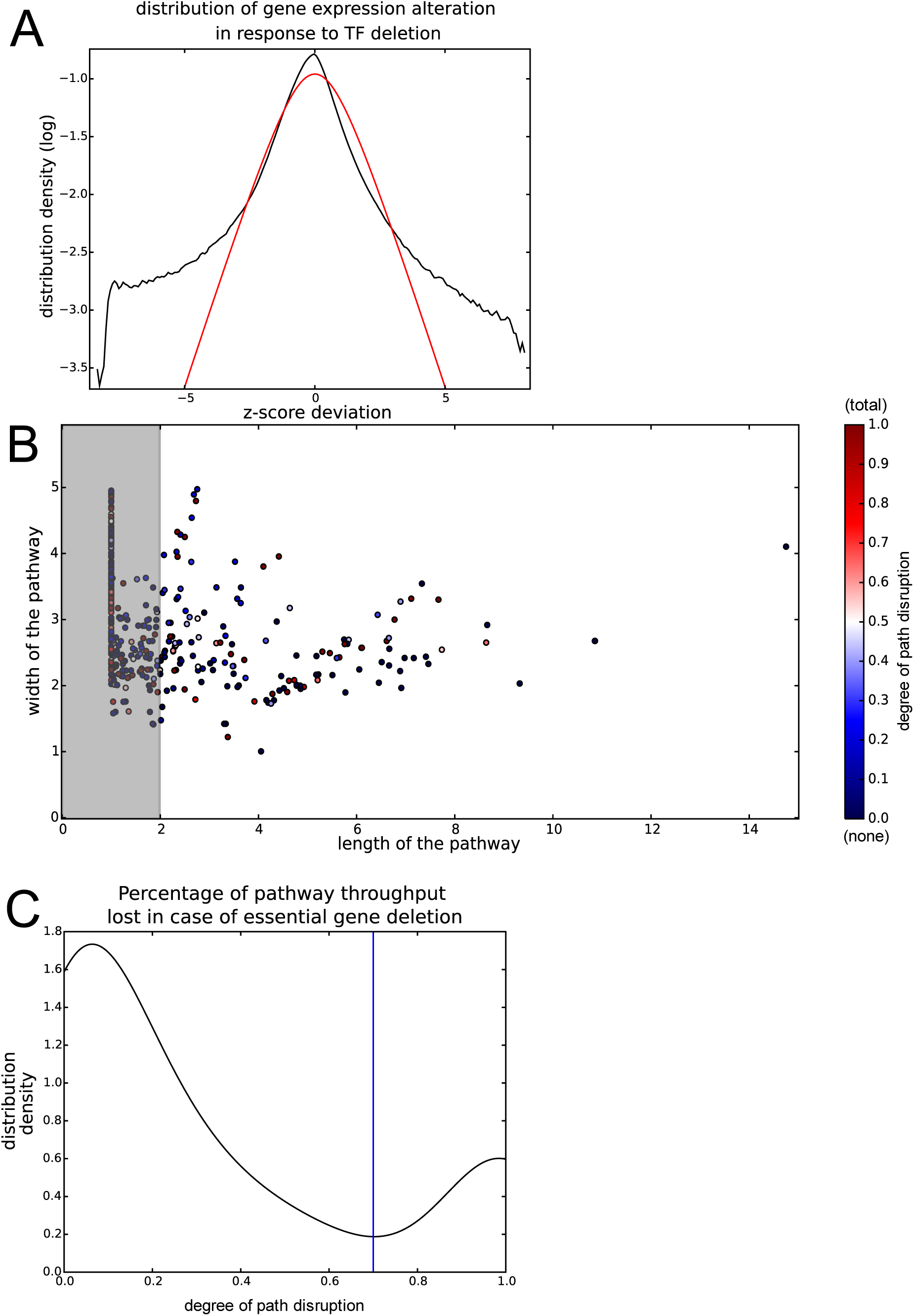
Data used to generate the simulated networks. (A) Black - distribution of TF deletion effects on gene product abundance in yeast, all 250 TF combined (Z-scores). Red - normal distribution centered around 0. (B) Correlation of non-trivial random pathways lengths and widths. Grey area designates width/length correlations discarded as likely corresponding to a tight hub (pathway length below 2). Color codes for the degree of pathway disruption by the deletion of an essential gene found in the path, whether the gene essentiality is associated to pathway function or not (see (23)) (C) Distribution of non-trivial random pathways disruptions upon essential genes deletion. Blue line - chosen threshold for critical network failure (∊) at 0.7.

Second, we estimate the length and width of pathways in yeast by using published protein-protein interaction (PPI) data (*24, 25*), as well as the manually curated Reactome database of biological reactions (*23, 26*). We define a *random pathway* as a set of paths in PPI and Reactome network that links two randomly chosen proteins, limited to most prominent paths in the entire network connecting them. Given that the master graph, combining all the links from PPI and Reactome, contains all the pathways, the pathways sampled in this way account for pathway overlap and interconnectedness (*23*). We obtain the distribution of pathway lengths (Supplementary Figure 1B) and widths (Supplementary Figure 1C), which exhibits a correlation (Figure 2B).

Finally, we retrieve a plausible value for *∊*, the maximal amount of perturbation of a pathway that is critical for organism survival. In order to calculate this value, we used the same dataset of PPI and Reactome, and for each random pathway, we evaluated how many paths were passing through a gene known to be essential (*11*). We obtained a two-peak distribution, as shown in Figure 2C. One peak is close to 0 and corresponds to no pathway disruption, representing the case where the gene was not playing an important role in the pathway and was present only by chance. Its essentiality may reside in a different pathway, whose genetic support partially overlaps with the pathway at hand. The second peak is centered around 1 and corresponds to a total pathway disruption upon gene deletion, meaning that this pathway depends critically on that gene. We see the transition from one distribution to another at around 0.7. There is no correlation between the importance of an essential gene for a random pathway and pathway shape, as shown in Figure 2B.

After retrieving the statistical graph parameters from experimental data, we used Equation 1 to estimate the number of essential genes. According to the estimated value for *∊* (*0.7*), essential genes would correspond to the genes for which deletion results in pathway throughput falling below 30% of its normal value. This normal value is computed by a simulation on a network representing a pathway assumed to be crucial for organism survival (*23*). The computation consists of a batch of feed-forward layered architectures for which length, width, and weights were drawn from experimentally obtained distributions.

To retrieve the statistics of essential gene abundance, at each round of simulation, we sample the essentiality statistics on 100 pathways. We expect the variation from these simulations to be representative of the variation in the fraction of essential genes that we would likely to find in organisms with a given statistical properties of the network. To get a sufficiently precise estimate of essential genes abundance, we performed 1000 rounds of simulations, which allowed us to calculate the distribution of essential genes abundance (Figure 3A, black), summarized by mean and standard deviation (Figure 3A, red). Our model’s 95% prediction interval for essential gene abundance is 9.2%-15.7% with a mean value at 12.5% (Figure 3A). The experimental value for the yeast essential genes is 16.9%. This value is not within the 95% prediction interval of our model but given the model’s simplicity and usage of raw experimental data, the experimentally measured value is still close.

**Fig. 3.**
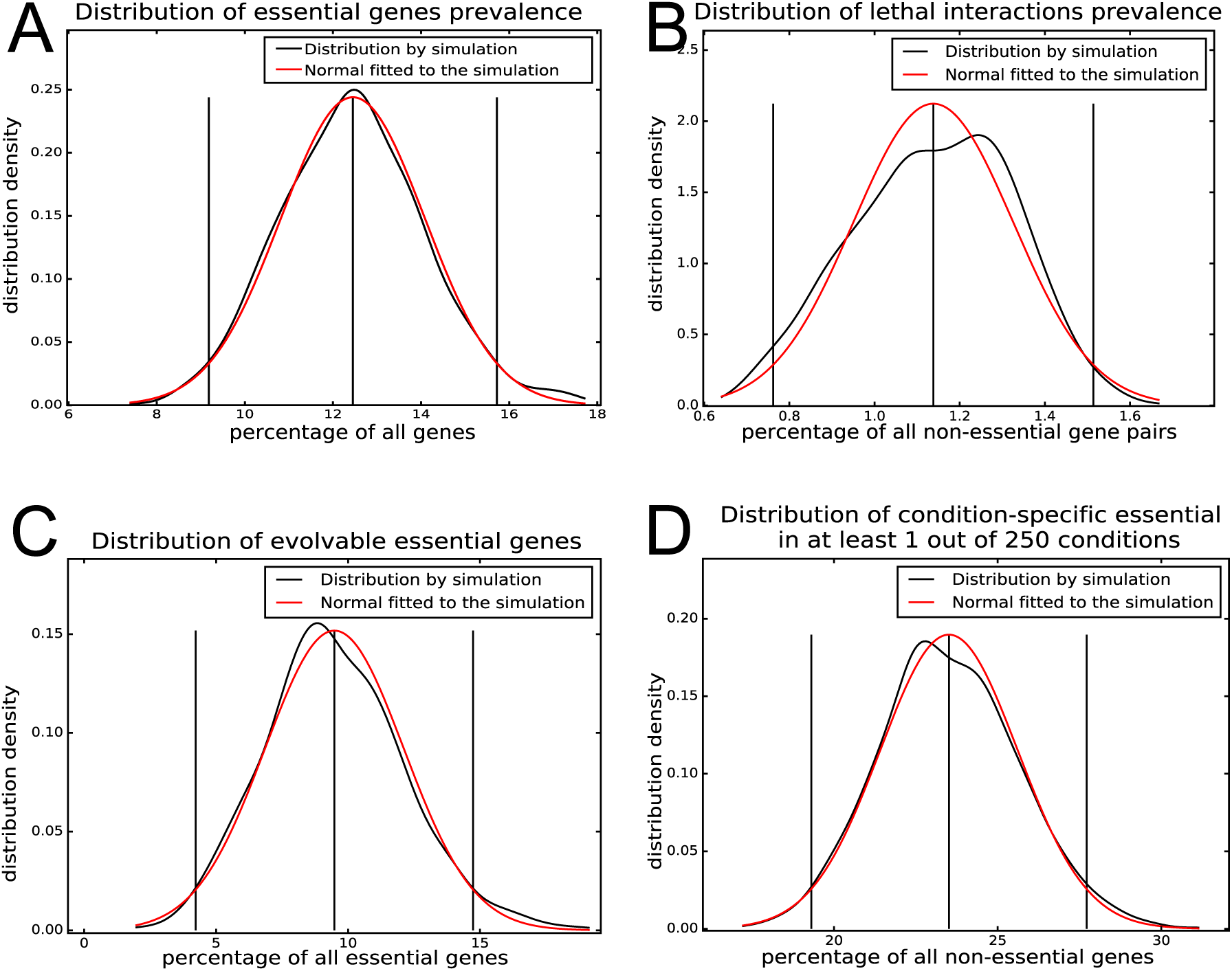
Results of simulation based on the model. (A) Estimates of the essential gene abundance according to our model. In black - distribution of essential genes abundances obtained by simulation, in red - Gaussian fitted to mean and standard deviation of the simulation data. Black vertical lines are mean and 95% prediction interval. (B) Estimates of the lethal interaction among non-essential genes in our model. In black - distribution of essential genes abundances obtained by simulation, in red - Gaussian fitted to mean and standard deviation of the simulation data. Black vertical lines are mean and 95% prediction interval. (C) Estimates of the abundance of evolvable essential genes among essential genes according to our model. In black - distribution of essential genes abundances obtained by simulation, in red - Gaussian fitted to mean and standard deviation of the simulation data. Black vertical lines are mean and 95% prediction interval. (D) Estimates of the abundance of genes that become essential in at least one of 250 stress conditions.

In addition to essential genes, BOWN also allows prediction of synthetically lethal (SL) interactions between non-essential genes. SL interactions within our model are co-deletions of non-critical nodes that would lead the error over the pathway to exceed 70%. Using the same notations as for Equation 1, genes *k*_1_ and *k*_2_ (*k*_1_ ≠ *k*_2_) in layers and *l*_1_ (not necessarily different layers) are SL when we have a similar condition as in Equation 1, with 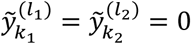 representing the simultaneous deletion of both genes.

Estimating the prevalence of SL interactions requires additional information compared to essential genes abundance estimation. While BOWN can only estimate the SL of interactions within the same pathway network, experimental data (*27*) provides SL interaction prevalence among all non-essential genes. We therefore need to account for the genes outside the pathway network at hand. Since there are about 17 independent pathway clusters in yeast (*27*), we add 16 times the number of genes in the pathway network as non-lethal interactions when computing the final statistic, to account the genes in similar non-overlapping networks. This addition corresponds to the intuition that to have a SL interaction, two genes need to be involved in the same physiological process, and that we expect that two randomly selected genes only have a 1/17 chance to belong to the same pathway. This gives predicted 95% prediction interval of 0.76%-1.5% with the mean value at 1.1% for the proportion of the interactions between non-essential genes being SL (Figure 3B). Once again, while not within the 95% prediction interval, this is still close to the experimental data suggesting a 1.62% prevalence of SL interactions in yeast (*27*).

BOWN can also be tested by attempting to estimate evolvable essential genes abundance. Evolvable essential genes are defined as genes that are essential in an unperturbed genome, but whose lethality upon deletion can be overcome through large scale alteration of gene stoichiometry such as through aneuploidy (*4*).

In order to capture the effect of aneuploidy on pathway networks, we estimated the perturbation of protein abundance from published proteomics data from aneuploid yeast (*28*). For each protein, we calculated the ratio of its abundance in a specific aneuploidy relative to its abundance in the diploid population. We then pooled all ratios from different aneuploids into a single distribution (Supplementary Figure 2A). We simulate aneuploidy by altering nodes activation levels by the factors drawn from that distribution. For each node considered essential without perturbation, we assessed whether the node deletion in the perturbed network would lead to the network throughput falling below the critical level. Aneuploidy generates a stress on the organism by itself, even in the normal conditions (*28*), which we reflect by increasing *∊* from 0.3 to 0.4, meaning that a smaller error on the output function could be fatal. This correction is based on previously reported growth deficiency of aneuploids of approximately 11%-13% compared to euploid cells (Pavelka et al. 2010), which we interpret as an ^~^10% lower tolerance to pathway disruption. BOWN estimates evolvable essential gene abundance to be 9.48% + 3.4% (mean ± SD), close to the 9% reported previously in experimental studies (*4*), which falls in the 95% prediction interval of our model (Figure 3C).

Another prediction of BOWN is that the distribution of essential genes will shift, not only as we alter the structure or average node activation of the biomolecular network, for instance with aneuploidy, but also as we vary the inputs of the network, corresponding to environmental changes. We randomly perturbed the inputs for the first layer in our simulations, changing them to values that were sampled randomly from a uniform distribution ranging from 50% to 150% of the values associated with the rich medium. To simulate different conditions, we chose 250 different input samples for each pathway network - we stopped at 250, since beyond 250, the prevalence of conditional essential genes does not change (cf. Supplementary Figure 2 B). Every gene whose deletion would lead the perturbed network output to drop below *30%* of its original value would be classified as a conditional essential gene in at least one condition. Our model predicts the distribution of conditional essential genes in at least one of the 250 conditions to be 23.5 ± 2.1% (Figure 3D). This prediction is consistent with experimental data suggesting that almost all yeast gene deletions that exhibit no phenotype in the rich medium can result in fitness loss in a stressful condition (*29*). Unfortunately, while the prior work mentions that almost 100% of yeast genes are required for optimal growth in at least one of the conditions, the experimental statistics for genes that are essential in at least one of the environments are not available. This abundance of condition-essential genes is a prediction of our model that will need to be validated experimentally in the future.

Finally, to verify that those predictions were specific to yeast biomolecular network and not due to general properties of the model, we altered the distributions used to build the modeled networks by plausible alternatives, i.e. given by closest standard deviation to the observed values. First, we replaced the distribution of pathway lengths and widths by a uniform distribution ranging from the minimum to the maximum values found in empirical data (Supplementary Figure 3A). Second, we replaced the edge weights distribution by a Gaussian distribution (Figure 2A, red). Third, we replaced aneuploidy distributions by a log-normal distribution with parameters *μ = 0* and *σ =* 0.5 (Supplementary Figure 3B, red). Finally, we chose *∊* to be 0.5 instead of 0.3. Table 1 summarizes the effects of these replacements. Overall, this shows that the predictions are indeed specific to the yeast biomolecular network and sensitive to the parameters that were experimentally observed. It is crucial to note that none of the parameters were fitted to allow us to retrieve experimentally observed results.

**Table 1.** Model sensitivity to the network parameters. Perturbations of network parameters according to likely distributions lead to a strong deviation of statistics predicted by the model from the experimentally observed ones.

|  | essential % | SL interactions % | evolvable essential % |
| --- | --- | --- | --- |
| experimental observations | 16.9 | 1.62 | 9 |
| base model | $12.5 \pm 1.6$ | $1.1 \pm 0.18$ | $9.5 \pm 2.6$ |
| uniform pathways shape | $7.9 \pm 1.2$ | $0.35 \pm 0.01$ | $8.8 \pm 2.4$ |
| normal activations | $10.0 \pm 1.7$ | $1.1 \pm 0.20$ | $14.5 \pm 3.1$ |
| log-normal aneuploidy effect | $12.6 \pm 1.3$ | $1.1 \pm 0.17$ | $24.8 \pm 3.5$ |
| $\epsilon = 0.5$ | $29.0 \pm 2.5$ | $2.2 \pm 0.24$ | $27.5 \pm 3.0$ |

Another prediction of BOWN is that networks are made more robust by decreasing the kinetic factors of reactions (*23*) (Lemma 2). For instance, slower gene expression leads to more robust organisms. Interestingly, this conclusion is a generalization of an earlier observation involving a single pair of interlinked feedback loop (*30*). The previous work suggested that slow activation was critical for robustness and noise resistance, whereas our theory makes a similar prediction in a more general context. BOWN suggests that this observation is connected to a more general and well-known dilemma of robustness versus rapid response in distributed computing (*31*). In distributed computing, safety (robustness) usually requires a compromise on liveness (rapid response) and vice-versa (*15*).

More generally, BOWN can account for perturbations of networks that are more specific than aneuploidy. Mutations or modifications of interaction partner patterns, such as common in cancer (*32, 33*) (cite), or upon organism evolution (*34, 35*) (cite) can also be accounted for by BOWN. Unfortunately, we do not have the data to properly validate BOWN with regards to those applications which constitute a promising avenue of future exploration.

In summary, BOWN predicts the abundance of essential and evolvable essential genes as well as SL interactions between non-essential genes. The predicted statistics are close to experimentally observed values. The predictive power of our model is remarkable, given the level of abstraction of the layered networks in our model compared to real biological networks. The relative accuracy of our model suggests that the positions of physical entities inside the biological computation graph are more important than the specific functions those physical entities accomplish.

Our results argue for the validity of modeling of biomolecular networks as *weighted layered graphs with nonlinear nodes*, which allows prediction of phenotypes associated with gene mutations at the network scale based solely on the high-level statistical properties of the network. This last feature enables functional dissection of biomolecular networks without being limited by the current lack of detailed network description. This could make BOWN a useful approach for improving our understanding of complex genetic phenotypes and diseases.

## Acknowledgments

We would like to thank J. Zhu, A. Baryshnikova, ES Deeds and J. Bader for the constructive feedback on the manuscript.

## Author contributions

EE & AK – original draft, conceptualization, methodology & investigation, EE – formal analysis, AK – data curation, software, visualization, RG & RL – funding acquisition, project administration, supervision, review and editing.

## Data and materials availability

Code is publicly available at the following repositories: https://github.com/chiffa/GeneNetworkStructure, https://github.com/chiffa/BioFlow: data was collected from public sources cited in the article.

## List of Supplementary Materials

(*36, 37*) are only used in supplementary material:

## Footnotes

1 They are called *single* in the sense that the failure of any one is fatal, even if they are *numerous*.

