## Supplementary information, Supp. Figures, and Supp Tables for "Predicting complex genetic phenotypes using error propagation in weighted networks"

**This PDF file includes:**

Materials and Methods
Supplementary text
Figs. S1 to S3

**Materials and Methods**

Retrieval of network statistics in yeast

In order to retrieve network statistics in yeast, we used a BioFlow tool instance (code available at https://github.com/chiffa/BioFlow), initialized with a yeast biomolecular network. To summarize, the SwissProt Uniprot database (*37*) was a backbone. Interaction data sources were the Reactome database of reactions in yeast (*26*), the HiNT high-quality yeast interactome (*24*) and the BioGrid PPI database for yeast (*25*). To account for possible mismatch, proteins corresponding to different databases, instances of proteins in different databases were created. To account for the presence of repeated instances of proteins as well as difference in the confidence we would have into the PPI interactions that would have a different level of confidence, we assigned weight to edges of biomolecular network. Connections coming from reactions in Reactome as well as from HiNT were assigned a weight of 1. Connections coming from BioGrid were assigned a weight of 0.5. Finally, edges identifying instances of different proteins in different databases, based on Uniprot data, were assigned a weight of 100. All the edges were considered as undirected.

In order to compute the shape of pathways, a connex set of nodes was extracted from the graph, two nodes chosen randomly in it and information flow calculated between them. In order to calculate the information flow, the two proteins were treated as a pair of sink/sources of electric current and the entire connex part of network treated as a support for electric current flow, with edges treated as resistances, with resistance conductance proportional to the edge weight. The computation was performed by extracting a Laplacian of the network, removing the column/line pair corresponding to the sink and using *scipy.sparse.linalg.splu* function in order to solve the equation corresponding to the matrix Laplacian with current set to 1.

Following the solution, the total intensity of current through the nodes was calculated and all nodes routing less than 5% of total current were removed. The width of the resulting *random pathway network* was calculated as the inverse of the current passing in the choke-point node, defined as the node with the highest flow outside the sink and source. The width and voltage difference were then used in order to estimate the lengths of the random pathway network. These values were directly stored.

In order to estimate the critical disruption in biological networks and hence $\epsilon$, we projected the known essential genes in yeast network onto the real random pathway networks and calculated the fraction of current the essential genes were routing in that network. The genes present in the network by chance would have a low flow through them, in the vicinity of 0. The genes whose essentiality is related to a real pathway we happened to include in our random pathway would be routing a large amount of flow, in the vicinity of 1. Only the flow through the essential node with the highest flow rate in the network was considered.

The protocol implemented here is implemented in code, available from https://github.com/chiffa/BioFlow, specifically *bioflow.utils.usage_ex-average_path_length.py*. A total of 701 random pathways were computed, of which all the pathways with a length below 2 were dismissed as likely belonging to a hub neighborhood in the PPI interaction network

Simulation of functional networks

In order to simulate a functional pathway network, we built a rectangular feed-forward network, fully connected layer-to layer. The width and length of the network were retrieved from the yeast biological pathways as described in section 2 of supplementary materials, whereas the weights were retrieved from the TF deletion data as described in the main text. Since we are interested in the highest error possible upon deletion, we converted the weights $c_{i,j}, K_{i,j}$ to their absolute values and removed node non-linearity. In this network, a forward pass corresponds to the maximum information that can be transmitted through the network by each node (Figure 1E)

To calculate if a node was critical and whether the corresponding gene would be considered essential, we first performed a reference pass of the error network, calculating the final output. Then we deleted one by one every node in the intermediate layers and monitored the output of the maximum error propagation through the network. If that maximum throughput fell below 30% of the original value, we declared that node as critical, interpreted as corresponding to an essential gene. All non-essential and essential nodes were memorized upon this first pass.

A second pass was performed by co-deleting all combination of pairs of non-essential nodes (Figure 1F). Whenever the total throughput on the error network fell below 30%, a pair was declared as corresponding to a pair of lethally interacting genes.

Aneuploidy was represented by randomly multiplying the activations of the nodes in the network by values randomly selected in the set of fractions of protein abundances between aneuploid yeast strains and euploid yeast strains in prior publications, at each layer, at each step of a forward pass (Figure 1E). Nodes from the essential set were then deleted one by one and we checked if their deletion would still drop the throughput of the error network below 30%. If this was no longer the case, we declared those nodes as evolvable essential.

The environment perturbation effect was calculated by introducing a random perturbation sampled from the uniform distribution with support on $[0.5,1.5]$ to the first layer of node activities in the network (Figure 1E). The conditional essentiality was calculated by iteratively deleting all the non-essential genes and checking if the nodes throughput fell below the 30% once the inputs were perturbed.

The full code of the simulation is available from https://github.com/chiffa/GeneNetworkStructure and the networks simulation, along with effect calculation was performed by the *Network_builder.py* module.

**Supplementary Text**

Viewing a Genetic Network as a Distributed Computing System

A distributed computing system has two main types of components, processes and communication channels (*13*, *14*).

Processes:

Processes are physical entities representing the computing nodes of the model. Whether small molecules, proteins, RNA or fragments of DNA, they route the information through a biological network by changing their state through a biochemical reaction, be it the modification of the compartment localization, the complex participation [1] or the transformation into a different physical entity. We model their unreliability by either a direct failure (corresponding to the inactivation of the gene associated to the node or the disappearance of a component for its synthesis) or by a random value of the output [2] (corresponding for instance to the import/export from/to the environment or a mutation modifying the gene promoter unpredictably). We assume that failures of different genes are completely independent one from another.

Communication Channels:

The information is transmitted between the nodes through the functional modification of one node by the other, be it by reaction co-participation, complex formation, that changes the properties of the upstream node, or post-transnational modification. The chemical relationships between nodes are viewed as *channels* and are weighted by a coefficient $c$. Since reactions are locally limited by the reagents, diffusion limits the rate of a reaction, meaning that the transmission channel is limited with respect to the intensity of transmitted information. We represent this by bounding the weight coefficient $c$ of a channel.

Computations:

Together, the processes and the communication channels in genetic networks constitute a weighted, layered and directed graph of which we describe the computation in what follows.

In order to represent the dynamics of a process, we use a simplified variant of the Hill (*27*) equation. This is a commonly used equation that captures the kinetics of the biochemical reactions in the network.

$$y_i^{(l+1)} = \frac{1}{(\sum_{j=1}^{N_l} c_{ij}^{(l)} y_j^{(l)})^{n_{ij}} + 1} = \varphi_i\left(y_1^{(l)}, \dots, y_{N_l}^{(l)}\right) \qquad (1)$$

In Equation (1), $y_i^{(l)}$ denotes the concentration of the physical entity produced by the biochemical reaction in node $i$ of layer $l$. $N_l$ denotes the number of nodes in layer $(l)$. By convention, $l = 0$ refers to the sensor layer (inputs), so that $y_i^{(0)} = x_i$ denotes the concentration of the physical entity corresponding the sensory node, similarly, $l = L + 1$ refers to the output (the genetic function). The input nodes are considered as given by the environment and do not follow a Hill equation. In contrast, the output of the network follows a Hill equation and are thus bounded. The coefficient $c_{ij}^{(l)}$ is used to abstract the stoichiometric relationship between node $j$ of layer $(l)$ and node $i$ of layer $(l + 1)$. For the output node we use a single low index as there is a single node in layer $L + 1$. Finally, $n_{ij}^{(l)}$ is the same factor as in the usual Hill equation.

In Figure 1 we sketch our simplified view of a genetic network. As a first approximation, we view this network as a directed graph where the information flows from the input (sensory) side on the left (denoted by the input vector $X$, to the genetic function on the right (denoted by

---

[1] When two or more molecules are combined.

[2] Byzantine failures, as in the traditional distributed computing vocabulary.

$F(X)$). Without loss of generality, we neglect the looping, the self-inhibitory and self- excitatory effects.

The computation performed by the graph is given by the following equations:

$$F(X) = \varphi(\sum_{i=1}^{N_L} c_i^{(L+1)} y_i^{(L)}(X)) \;\; with \; y_j^{(l)} = \varphi(s_j^{(l)})(l \geq 1);\; y_j^{(0)}(X) = x_j \; and \; s_j^{(l)} = \sum_{i=1}^{N_{l-1}} c_{ji}^{(l)} y_i^{(l-1)} \qquad (2)$$

Results:

We establish a theoretical condition for a single node to be critical in weighted directed graph with sigmoidal units. We prove this condition using a theoretical assessment of error propagation in such graphs. We first introduce the following useful lemma:

**Lemma [lipschitz]:** The function $\varphi$ from the Hill equation is $n_{ij}^{(l)} c_{ij}^{(l)} (N_l c_i^{max(l)})^{n_{ij}^{(l)}}$-Lipschitz [3] with respect to variable $y_j^{(l)}$, $c_i^{max(l)}$ being the maximal weight of the incoming edges to node $i$ of layer $l$.

(Sketch) For notational convenience, we will omit the reference to the layer ($y_j = y_j^{(l)}, c_{ij} = c_{ij}^{(l)}$ and so on). Note that $\varphi(y_1' \dots, y_{N_l}) = \frac{1}{(\sum_{j=1}^{N_l} c_{ij} y_j)^{n_{ij}+1}} = \frac{1}{1+exp(n \cdot log(\sum_{j=1}^{N_l} c_{ij} y_j)}$. Taking the partial derivative w.r.t to variable $y_j$ we have $\frac{\partial \varphi}{\partial y_j} = - \frac{\frac{n_{ij} \cdot c_{ij}}{\sum_{j'=1}^{N_l} c_{ij'} y_{j'}} \cdot exp(n_{ij} \cdot log(\sum_{j'=1}^{N_l} c_{ij'} y_{j'})}{(1+exp(n_{ij} \cdot log(\sum_{j'=1}^{N_l} c_{ij'} y_{j'})))^2}$ of which the absolute value is bounded by $n_{ij}^{(l)} \cdot c_{ij}^{(l)} \cdot (N_l c_i^{max(l)})^{n_{ij}^{(l)}}$. Denote by $K_{ij}^{(l)} = n_{ij}^{(l)} \cdot c_{ij}^{(l)} \cdot (N_l c_i^{max(l)})^{n_{ij}^{(l)}}$ in all what follows. By the theorem of intermediate values, $\forall x, y \; \exists z \in (x, y)$ such that:

$$\frac{\varphi(x_1, x_2, \dots, x_{j-1}, x, \dots) - \varphi(x_1, x_2, \dots, x_{j-1}, y, \dots)}{x - y} = \frac{\partial \varphi}{\partial y_j}(x_1, x_2, \dots, x_{j-1}, z, \dots).$$

The Lipschitz inequality follows from $\frac{\partial \varphi}{\partial y_j}(x_1, x_2, \dots, x_{j-1}, z, \dots) \leq K_{ij}^{(l)}$. This inequality is tight since the first derivative of $\varphi$ reaches its upper bound ($\varphi$ in continuously differentiable taking values in compact domain).

As a consequence, we can derive a series of necessary conditions to estimate the number of essential genes, i.e genes whose failure is lethal.

**Proposition [sufficient]:** Following our notation, the number of essential genes is lower-bounded by

[3] Recall that a function (x1,…,xN)↦φ(x1,…,xN) is K-Lipschitz w.r.t variable xj if ∀(x,y)|φ(x1,x2,…,xj-1,x,…)-φ(x1,x2,…,xj-1,y,…)|≤K.|x-y|. We consider the Lipschitz coefficient as the smaller such K, this guarantees the tightness of our bounds and the quality of our estimators.

$$\left|\left\{(l,i) \in [1,L]\times[1,N_l] s.t \exists j \in [1,N_{l+1}]: c_{ij}^{(l+1)} K_{max}^{(l)} \left(\prod_{l'>l}^{L+1} N_{l'} c_{max}^{(l'+1)} K_{max}^{(l')}\right) > \epsilon\right\}\right| \quad (3)$$

where $|S|$ refers to the cardinality of a set $S$ and $\epsilon$ is the minimal lethal error at the output. When the network is optimal, this lower-bound is tight.

Proof (sketch): Proposition [sufficient] is proven by induction on $L$ (the number of layers). Initiation ($L = 1$) follows from the Lipschitz properties as detailed in (El Mhamdi and Guerraoui 2017). Induction step consists in assuming the bound valid up to a given integer $L$ and seeing the nodes of layer $L + 1$ as outputs of an $L$-layered network, then applying the $L = 1$ result on the output layer (the $(L + 2)$-th layer).

Detailed proof: we first need the following lemma on error propagation.

Let us denote by $F_{fail}$ the value of the metabolic function when a single node is failing. We keep referring to the nominal metabolic function (when no node is failing) by $F$.

**Lemma [aux]** (Error propagation from single fail): Let $X$ denote any vector of nutritional inputs. In a feed-forward sigmoidal graph, the failure of node $i$ in layer $l$ yields a propagated error through node $j$ in layer $l + 1$ to the output, bounded as follows:

$$\| F(X) - F_{fail}(X) \| \leq c_{ij}^{(l+1)} K_{max}^{(l+1)} \left(\prod_{l'>l+1}^{L+1} N_{l'} c_{max}^{(l'+1)} K_{max}^{(l')}\right) \quad (4)$$

This bound is tight.

We proceed by induction on the number of layers L.

**Initiation**. In the base case, L=1, let $X$ be any input vector, following equation (4), the error at the output when a neuron -denote it $k$ - in layer 1 fails is

$$\| F(X) - F_{fail}(X) \| = \| \varphi\left(\sum_{i=1}^{N_1} c_i^{(L+1)} y_i^{(L)}(X)\right) - \varphi\left(\sum_{i=1, i\neq k}^{N_1} c_i^{(L+1)} y_i^{(L)}(X)\right) \| \quad (5)$$

by the Liptschitz property of $\varphi$, and the definitions of $c_{max}^{(2)}$ and $K_{max}^{(2)}$ respectively as the maximal weight from layer 1 to the output (layer 2) and the maximal Lipschitz coefficient at layer 2, the right term is tightly upper bounded as follows:

$$\| \varphi\left(\sum_{i=1}^{N_1} c_i^{(2)} y_i^{(L)}(X)\right) - \varphi\left(\sum_{i=1, i\neq k}^{N_1} c_i^{(2)} y_i^{(L)}(X)\right) \| \leq c_k^{(2)} K_{max}^{(2)} \leq c_{max}^{(2)} K_{max}^{(2)} \quad (6)$$

, which is the desired upper bound.

**Heredity.** Let us now assume that the bound holds for any network of $l$ layers up to a given integer $L$. Consider a network of $L + 1$ layers where a single node failed. Recall that layer $L + 2$ denotes the output. We can see the $L + 1$-th layer as the output of a $L$-layer network, which in turn acts as a single layer, feeding the $L + 2$ layer.

- If the failing node is in layer $L + 1$, the single layer case proved above applies and the bound corresponds to the bound of Lemma [aux].

- If the failing node is in one the $L$ first layers. Following the inductive hypothesis, layer $L + 1$ is acting as single layer where the input vector is altered by at most $err$ such that

$$err = c_{ij}^{(l+1)} K_{max}^{(l+1)} (\prod_{l'>l+1}^{L+1} N_{l'} c_{max}^{(l'+1)} K_{max}^{(l')}),$$

by the Liptschitz property, this yields a propagated error of by at most $err.c_{max}^{(L+2)}K_{max}^{(L+2)}$ and we have

$$err.c_{max}^{(L+2)}K_{max}^{(L+2)} = c_{ij}^{(l+1)}K_{max}^{(l+1)}(\prod_{l\prime>l+1}^{L+2} N_{l\prime}c_{max}^{(l\prime+1)}K_{max}^{(l\prime)})$$

which is the desired bound for a $L+1$ layer network. By induction, the bound is proven for every integer $L$.

From lemma [aux], it follows that any node in the set $S$ of nodes defined by

$$S = \{(l,i) \in [1,L] \times [1,N_l] s.t \exists j \in [1,N_{l+1}]: c_{ij}^{(l+1)}K_{max}^{(l)}(\prod_{l\prime>l}^{L+1} N_{l\prime}c_{max}^{(l\prime+1)}K_{max}^{(l\prime)}) > \epsilon\}$$

will yield an error in the output larger than $\epsilon$, which is the minimal lethal error at the output. The set $S$ is therefore included in the set of lethal nodes. Therefore, the cardinal of $S$, $|S|$, is a lower bound on the number of essential nodes. Following the tightness of the upper bound in Lemma [aux], $|S|$ is a tight lower bound on the number of essential genes. QED.

Simply stated, Proposition [sufficient] translates to the following: a gene $i$ in layer $l$ is essential if there exists a gene $j$ in layer $l+1$ such that the forward propagated error through gene $j$ because of the knock-out of gene $i$ is lethal (larger than $\epsilon$), where $\epsilon$ is a lethal error on the metabolic function. The set of which we compute the cardinality is the set of such genes. The quality of Proposition [sufficient] as an estimator of the number of essential genes is due to the tightness of the lower-bound. With this tightness consideration in mind, the simplified model will look at nodes performing in the linear regime (maximal slope of the function $\varphi$).

In practice, experimental data on gene deletion can only provide an estimation for the product $c_{ij}^{(l+1)}K_{max}^{(l)}$ and not for the individual values of $c_{ij}^{l}$ and $K_{max}^{l}$ separately. Deconvoluting $c$ from $K$ in that product requires information on every single interaction, which is prohibitively time-consuming for experimental genetics. The multiplicative nature of our essential genes formula only requires a sampling of the $c.K$ products, not the individual values of $c$ and $K$. For now, we will see this product as the weight between nodes.

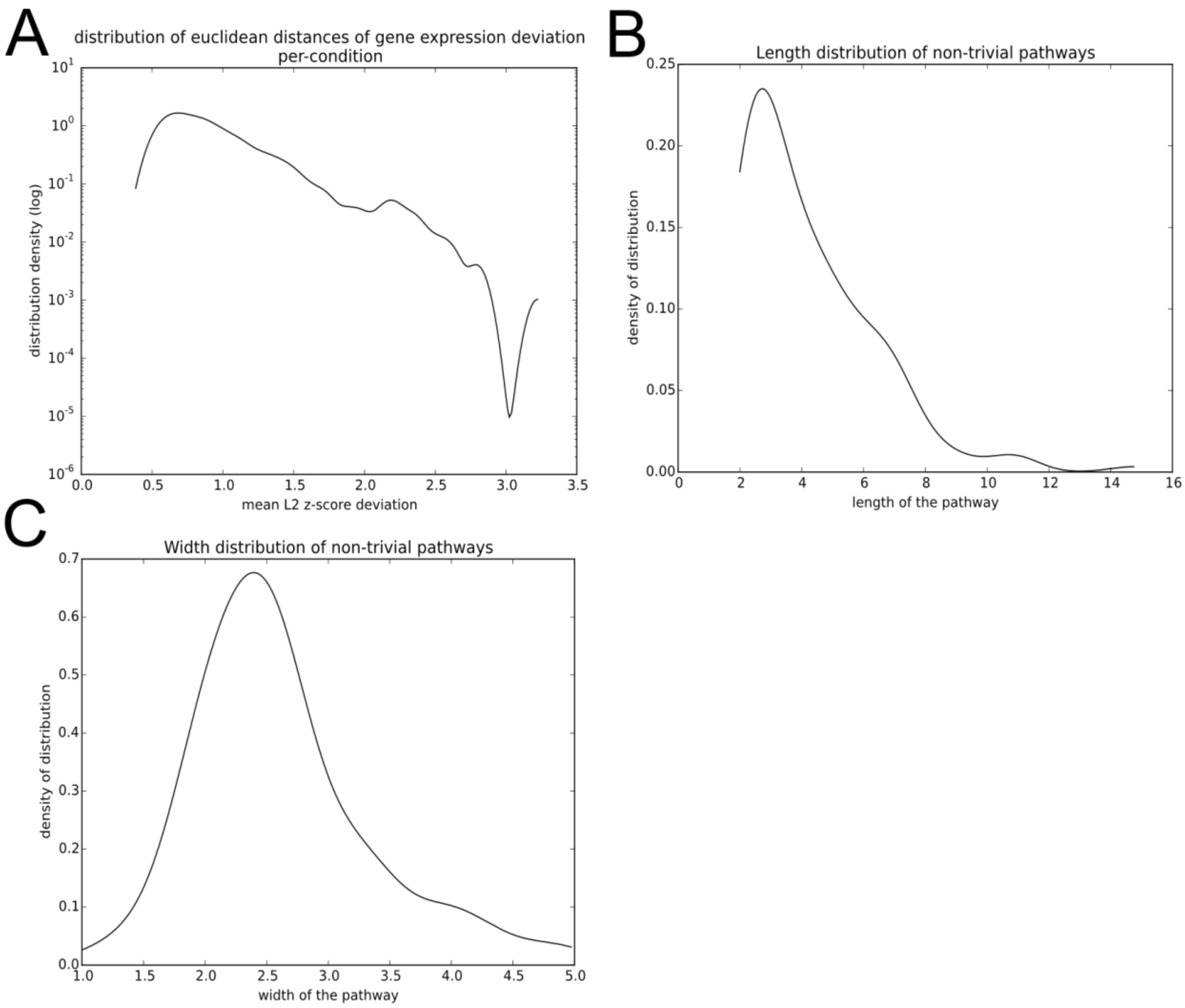


**Fig. S1.**

Additional data on the source data distributions. (A) Distribution of the average Euclidean deviation of gene expression upon TF deletions, for each condition, across all the genes. (B) Distribution of non-trivial random pathway lengths. (C) Distribution of non-trivial random pathways widths

A

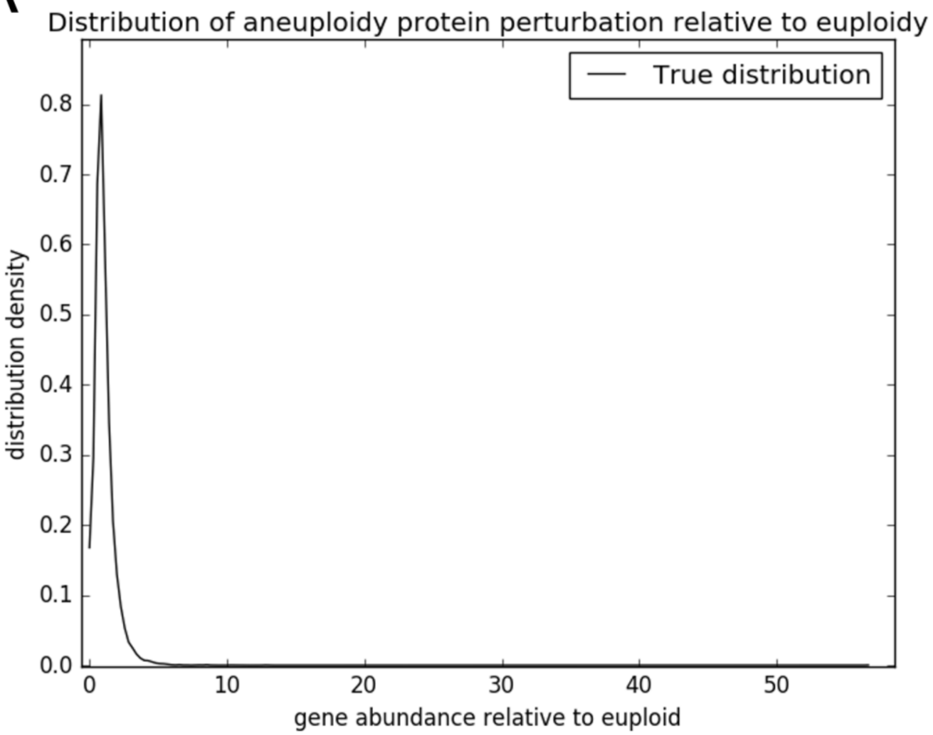
Distribution of aneuploidy protein perturbation relative to euploidy
True distribution
distribution density
0.0
0.1
0.2
0.3
0.4
0.5
0.6
0.7
0.8
0
10
20
30
40
50
gene abundance relative to euploid

B

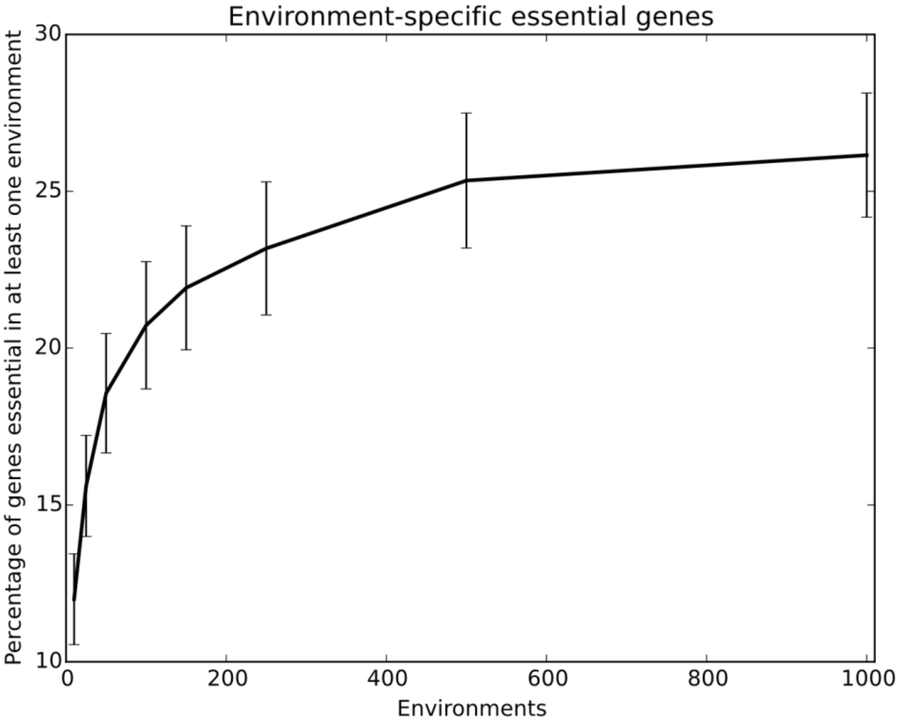
Environment-specific essential genes
Percentage of genes essential in at least one environment
10
15
20
25
30
0
200
400
600
800
1000
Environments

**Fig. S2.**

Additional data for essential genes in aneuploids and environment sets. (A) Distribution of protein dosage perturbation in a sample of aneuploid yeast relative to an isogenic euploid sample. (B) Prevalence of the genes that are essential in at least one of N environments, where N has been plotted on the x axis whereas prevalence – on the y axis. 250 represents good compromise between achieving an accurate evaluation of the condition-essential gene prevalence and computational load

A

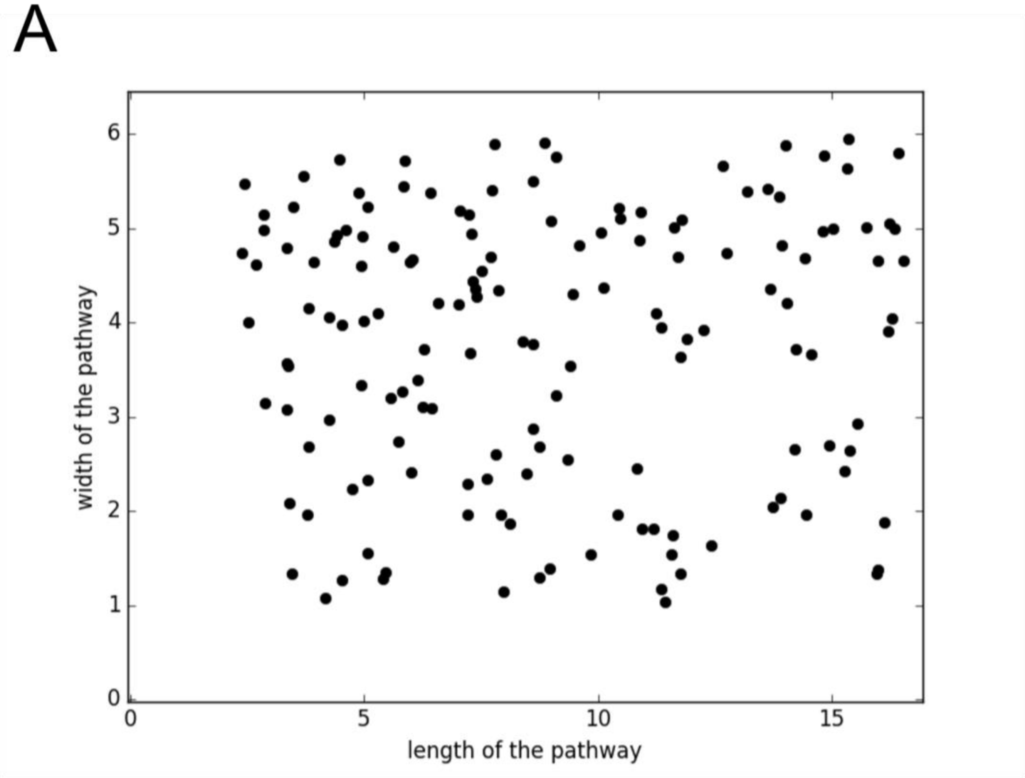
width of the pathway
length of the pathway
0
1
2
3
4
5
6
0
5
10
15

B

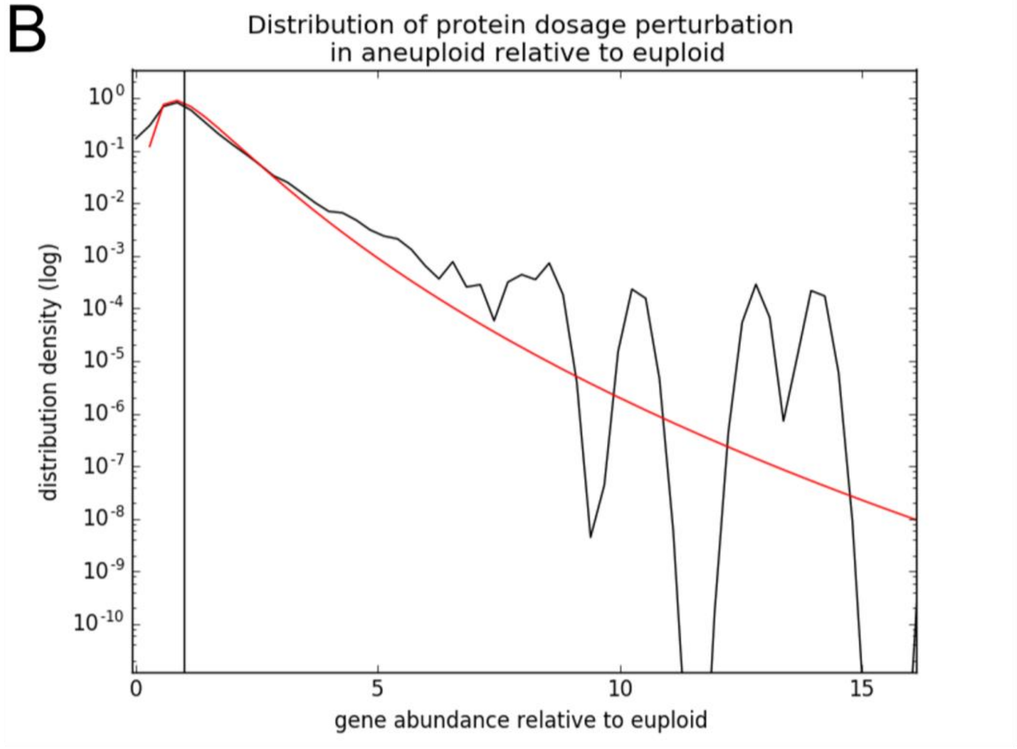
Distribution of protein dosage perturbation
in aneuploid relative to euploid
distribution density (log)
gene abundance relative to euploid
$10^{0}$
$10^{-1}$
$10^{-2}$
$10^{-3}$
$10^{-4}$
$10^{-5}$
$10^{-6}$
$10^{-7}$
$10^{-8}$
$10^{-9}$
$10^{-10}$
0
5
10
15

**Fig. S3.**

Additional distribution characterization for blank distributions validations. (A) Random pathway networks widths/lengths. (B) Comparison of protein abundance in aneuploid relative to euploid in experimental data (black) compared to log-normal distribution with shape parameters $\mu=0$ $\sigma=0.5$ (red). The black vertical line is at 1.0.
